# Hybrid Tetramers of Wild-Type and Amyloidogenic H88R Transthyretin Challenge the Monomer Paradigm

**DOI:** 10.64898/2026.07.13.738148

**Authors:** Zsolt Fazekas, István Bódy, Helga Tar-Pálfi, Dávid Papp, Dávid Virág, Lilla Turiák, Zoltán Pozsonyi, Dóra K. Menyhárd, András Perczel

## Abstract

Hereditary transthyretin amyloidosis (ATTRv) is driven by mutations that destabilize the native tetrameric transthyretin (TTR), promoting monomer formation and amyloid aggregation. The H88R variant has been considered fully monomeric, yet its behavior in heterozygous patients has remained unclear. Here, we demonstrate that H88R TTR can form hybrid tetramers with wild-type chains, both in vitro and in patient sera. We considered all possible tetramers, along with various trimer and dimer constructions, and found that the stability of hybrid tetramers containing one or two mutant chains is not significantly reduced in comparison with of the wild-type tetramer, suggesting a practically unhindered entry for H88R TTR monomers into such hybrid tetrameric assemblies, facilitating their secretion. Mass spectrometry confirms incorporation of H88R chains into tetramers in vitro, and show that the H88R TTR variant is present in the serum of carriers albeit in low concentration. We propose that the scarcity of this variant is the result of its retention in the endoplasmic reticulum and present a model for the association of H88R TTR with the endoplasmic reticule chaperone Binding immunoglobulin Protein (BiP). These findings revise the current monomeric view of H88R TTR, with direct implications for the efficacy of tetramer-stabilizing therapeutics. Our results highlight a delicate balance between cellular retention and hybridization, informing mechanistic understanding and treatment strategies in heterozygous ATTRv patients.

## Introduction

Proteins occasionally misfold, either prompted by inherent instability, or changes in circumstance or interaction partners. These misfolded states are typically unable to perform the function of the native protein, and are often more difficult to degrade through standard cellular housekeeping processes. These two properties cause the buildup of molecular dead weight in cells, taking useful space away from vital biochemical processes, which results in the impairment of cellular functions and, subsequently, cellular death. Amyloidosis is a disease, where the misfolded proteins arrange themselves in mostly β-sheet containing, regularly repeating layers, which build up the insoluble amyloid fibril macrostructure. The compactly packed layers lend a high stability to the fibril, making it resilient against the standard cellular degradation pathways. A wild variety of proteins are able to cause amyloidosis, such as immunoglobulin light and heavy chains^1–3^, the amyloid β-peptide^4,5^, α-synuclein^6^, prion proteins^7^, the serum amyloid A protein^8^, or the transthyretin (TTR) protein.

TTR is a 147 residue long protein (UniProt code: P02766), which, among the thyroxine-binding globulin and albumin, is responsible for the transport of thyroxine (T_4_) and (in complex with retinol binding protein) retinol, both in blood serum and the cerebrospinal fluid (CSF). Additionally, it has been suggested that TTR has important roles in neurological processes, such as neuritogenesis^9,10^, memory formation^11^, progression of Alzheimer’s disease^12,13^, or modulation of the noradrenergic system^14^. Structurally, in its native form TTR arranges itself into fold-protecting homotetramers (**Figure 1**, **Panel A**), where each chain contacts all others through three, qualitatively different interfaces (i.e. there are 6 interfaces in total in a tetramer). The monomeric units contain 8 β-strands and a short helix in a β-sandwich fold. Here, we refer to these interfaces as the β-edge (βE), β-cross (βC) and loop-loop (lL) interfaces (**Figure 1**, **Panels B** and **C**). The βE interface is a backbone-backbone antiparallel β-edge hydrogen-bonding network between two ^115^SYSTT^119^ sequences (residue numbering is according to the PDB ID 2QGB^15^). The βC interface is constructed from a pair of β-sheet triplets (regions K^15^-P^24^, T^106^-L^111^ and S^117^-N^124^) facing each other along the amino acid side chains. The hydrophobic amino acids in these regions facilitate the formation of the thyroxine binding cavity^16^, of which two is present in the full tetramer. The size of the cavity, when empty, allows the incorporation of structural water molecules, as can be seen in the crystal structure 2QGB. Finally, the lL interface is a loosely bound connection between two loop triplets (regions ^18^DAVRGS^23^, ^84^ISP^86^ and ^112^SPYS^115^). Using the PDBePISA webtool^17^, it can be determined that the βE, βC and lL interface contact areas measure 886 Å^2^, 367 Å^2^ and 329 Å^2^, respectively.

**Figure 1.**
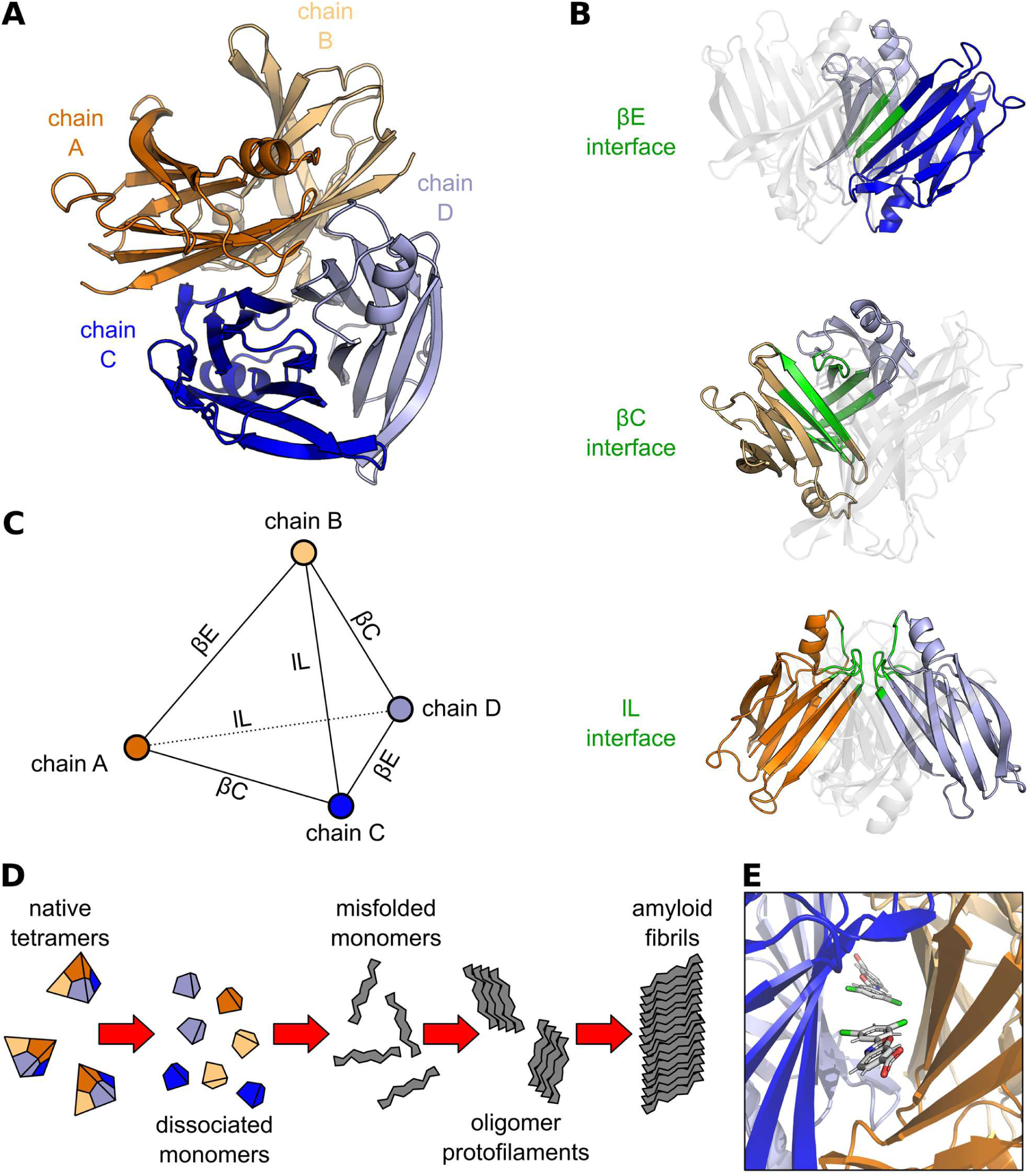
Panel **A** shows the tetrameric structure of the transthyretin complex. The different chains are indicated by four different colors. Panel **B** highlights the three possible interfaces between individual chains. Residues participating in the formation of the corresponding interface are highlighted with green. Panel **C** is a schematic of the relationship between the chains and the interfaces between them. The complex resembles a tetrahedral topology. Panel **D** is a schematic of the amyloid formation process. The transthyretin tetramers dissociate to monomers, lose their native fold and start associating into protofilaments. The protofilaments subsequently create the amyloid macrostructures. Panel **E** shows the structure of the tetramer stabilizing drug tafamidis in its binding groove.

TTR causes amyloid formation either through the aggregation of wild-type (WT) chains (wild-type transthyretin amyloidosis, ATTRwt) or through the aggregation of a mutant form (hereditary transthyretin amyloidosis, or variant ATTR, denoted by ATTRv) (**Figure 1**, **Panel D**). The onset time and prognosis of these diseases highly depend on the exact form of the aggregating TTR protein. ATTRwt usually develops in older age groups, mainly affecting men^18^. The amyloid depositions in ATTRwt accumulate mainly in the myocardium, leading to increased ventricular wall thickness and subsequent heart failure with preserved ejection fraction^19,20^. Carpal tunnel syndrome is also common in ATTRwt^21^. Similarly, ATTRv can manifest in a neurological or in a cardiac phenotype, but with markedly earlier onset. In contrast to ATTRwt, ATTRv is a familiar disease, transmitted in an autosomal dominant manner with incomplete penetrance. Depending on the underlying mutation, it can cause infiltrative cardiomyopathy and/or length-dependent sensory motor polyneuropathy, while also affecting other organs (eyes, kidneys and the gastrointestinal tract)^22^. With at least 200 single-point mutations considered pathogenic^23^, the most common ones in Europe include V30M, F64L, E89Q and T49A^24^. These mutations usually differ in tetramer destabilization tendency, disease manifestation, time of symptomatic onset and penetrance.

Current treatment options for ATTRv include the administration of tetramer stabilizing drugs (such as tafamidis, acoramidis and diflunisal), liver transplantation and gene silencing strategies (such as using an antisense oligonucleotide (inotersen, eplontersen) or a small interfering RNA (siRNA, patisiran, vutrisiran))^22^. Gene silencing therapies are administered either intravenously or subcutaneously, while stabilizers are oral medications. Due to the availability of effective pharmacological therapies and the invasive nature of the procedure, liver transplantation is now rarely performed. Currently, transthyretin stabilizers represent the most widely used specific treatment worldwide. Out of the available stabilizer drugs, tafamidis and acoramidis are certified medicinal products against ATTR cardiomyopathy, while the efficacy of diflunisal is yet to be assessed and is not yet certified. Furthermore, due to the previously mentioned neuroprotective effects of TTR, the use of gene silencing options might have negative neurological consequences.

Tetramer stabilizing medications, such as tafamidis, acoramidis and diflunisal, rely on the presence of the thyroxine binding cavity on the βC interface, where they can bind and mediate the inter-chain interactions of the complex (**Figure 1**, **Panel E**)^25–27^. All three of these molecules are benzoic-acid derivatives. The carboxylate moiety of tafamidis and acoramidis faces away from the core of the TTR complex towards the surrounding solvent, and is held in place by an ionic interaction with K^15^. The para- and meta-substituents of these molecules plunge into a hydrophobic cavity created by L^17^, T^106^, A^108^, L^110^, T^119^ and V^121^. In contrast, diflunisal is flipped in a way that the carboxylate faces the core and creates a hydrogen-bond with S^117^. The K^15^ residue is in a polar interaction with one of the fluoride substituents on the other aromatic ring. As the cavity is formed by residues from neighboring monomers, it is a sound assumption that tetramer stabilization can only occur if the tetramer is able to assemble by itself.

Position H^88^ in the TTR protein plays an important structural role. It has been shown that the unfolding propensity can be fine-tuned by selecting different TTR mutations at the H^88^ position^28,29^. This residue is located between the EF-helix (T^75^-G^83^) and the F β-strand. It mediates interactions on the βE interface directly and also through water molecules, but due to its proximity to other chains, it can also allosterically alter the stability and behavior of βC and lL interfaces. The only documented and medically relevant mutation at this position is H88R^23^, which is mostly prevalent in Austria^30^ and Hungary^31^, along with a few linked cases in Sweden^32^. This mutation causes the total disruption of the tetramer, as the bulky arginine side chain sterically hinders the formation of the βE interface, which would be the first step in the complexation reaction. This way, the H88R mutant TTR protein is known to form only monomers *in vitro*.

In here, we investigate the possibility of hybrid WT – H88R heterotetramer formation. Since small molecular treatment options focus on the stabilization of the already present TTR complex, it is crucial to investigate whether they are effective in the case of H88R mutant heterozygous patients. This stabilization can only happen, if the mutant TTR chains are co-located and are able to form mixed complexes with WT TTR chains. This must mean that the hybrid interface energies should be more beneficial or comparable to the WT – WT interface energies. Examples for such hybrid complexes are known for other TTR mutants (L55P, V30M, F87A, T119M)^33,34^. In our previous work, we pointed out the structural connection between H88R and other dissociation-prone variants, such as D18G, A25T and Y114H^35^. In those mutations, it has been already shown that patient serum TTR levels are noticeably below the physiological range. Especially, variant forms are not detectable or have a much lower concentration relative to the WT TTR present in the serum^36–38^. It was hypothesized, and later shown, that the Endoplasmic Reticule Associated Degradation (ERAD) hinders the secretion D18G and A25T *in vitro*^39^. We argued that H88R might follow the same path, based on the low serum TTR levels of H88R carrier patients as indirect evidence. Proving the feasibility of H88R-WT hybridization also opens a possible route for H88R to evade ERAD: it can possibly “hijack” the WT tetrameric complexes and get secreted in the form of hybrid tetramers. We performed *in silico* and *in vitro* studies to confirm the existence of H88R hybrid tetramers, which gives new insights into the treatment of H88R carrier patients.

## Materials and Methods

### Monte Carlo multiple minimum (MCMM) conformation search of the BiP-TTR complexes

1500 step MCMM-LMOD conformational search^40^ as implemented in the Schrödinger Modeling Suite (Schrödinger Release 2025-4: MacroModel, Schrödinger, LLC, New York, NY, 2025.) Each step of the search involved the random rotation (between 0-180°) of a randomly selected subset all torsion angles and random extent of relative translation (between 1-5 Å) and rotation (between 0-180°) of the two components (BiP and TTR), interspersed with LMOD steps (with a probability of 0.5), followed by energy minimization. Starting structure was built using PDB entries 9H0C and 3IUC. Torsional rotations were applied for TTR residues 72-91, but energy minimization was carried out, beside this segment, for BiP residues 197-220, 236-245 and 393-406 (comprising the binding site), keeping the rest of the model frozen. The global minimum energy complex structures thus obtained were subjected to MM/GBSA calculation of the Prime module of the Schrödinger Suite (Schrödinger Release 2025-4: Prime, Schrödinger, LLC, New York, NY, 2025.) using the OPLS4 forcefield^41^ to estimate the binding energy.

### Molecular Dynamics Simulations

All simulations were started from the structure with a PDB ID of 2QGB^15^. H88R mutations were introduced using the *PyMOL Molecular Graphics System, Version 3.0.4 Schrödinger, LLC*. Simulations were done using the GROMACS 2022.3^42,43^ software package with the AMBER99SB-ILDNP*^44^ force field and OPC water model^45^. The NaCl concentration was set to 150 mM. After system preparation, a four-step energy minimization and a four-step NVT equilibration was performed. In both of these, the protein atoms were positionally constrained to their initial positions with force constants of 1000, 500, 100 and 0 kJ×mol^-1^×nm^-2^. After the unconstrained NVT equilibration, a 100 ps long NPT equilibration took place. All production MD runs lasted 1000 ns at a temperature of 310 K (using the *V-rescale* thermostat in GROMACS) and 1 bar (using the *Parrinello-Rahman* barostat in GROMACS). The protein and the rest of the system were separately coupled to the thermostat. The step size was set to 2 fs and the leap-frog integrator was used.

### Trajectory Analysis Methods

The convergence point of the trajectories were assessed manually based on the root mean squared deviation (RMSD) (see **Supplementary Figures 1**, **2** and **3**) time dependence graphs. The maximum amount of time a tetramer trajectory took until convergence was 200 ns and this was used as a starting time for all analysis and for all trajectories.

MM/GBSA analyses were done using the gmx_MMPBSA 1.6.3 software^46^ based on MMPBSA.py 16.0 and AmberTools 20^47^. The single trajectory approach was used, i.e. the ligand, receptor and complex contributions were all calculated from a single trajectory file. In these runs, the leaprc.ff99SBildn force field was employed. In total, 8001 equally spaced frames per trajectory were used for enthalpy calculations, and 10 equally spaced frames per trajectory were used for normal mode analysis (NMA). NMA runs were done in order to get the entropy contributions. For the generalized Born calculations the GB-OBC1 model^48^ was used. For every trajectory, the TTR tetramer was dissociated and associated into dimer-pairs along all three interfaces, and the association Gibbs-free energies at 298 K are reported.

Residue – residue contacts were analyzed using RING 3.0.0^49^, while contact statistics and visualizations were extracted as previously described^50^, using in-house Python 3 scripts. RING analysis have been performed starting from 200 ns for every 500 ps of the trajectories (1601 frames, in total).

Principal component analysis was done using the *gmx covar* and *gmx anaeig* commands. Only the backbone and Cβ atoms were included in the covariance and subsequent eigenvector calculations. Rigid body segmentation^51^ was performed on the Cα atoms with a cutoff of 0.175 nm and with a Δ*t* of 100 ps.

### Notation of Complexes and Association Reactions

We simulated all possible TTR associations: dimers, trimers and tetramers. Tetramers are assumed to form regular tetrahedra, meaning that all chains are equivalent.

Tetramers are denoted by tuples of (4, *N*x, I), where *N* is the number of mutant chains in the complex and I is the relative position of the mutations if *N* = 2 (otherwise I is not indicated). For example, (4, 0x) is the completely WT homotetrameric TTR, (4, 1x) is a tetramer containing a single H88R mutant chain, and (4, 2x, βE) is a tetramer containing two H88R mutant chains, which are connected through a βE interface. As all tetramers were investigated in dimer-association reactions, an association reaction can be denoted by tuple pairs (I_dimer_)(4, *N*x, I_mut_), where I_dimer_ indicates how the dimers are assembled, and the rest of the notation indicates the target tetramer. For example, (βE)(4, 3x) denotes the association of two βE dimers to a 1:3 = WT:H88R tetramer, while (lL)(4, 2x, βC) denotes the association of two hybrid lL dimers to a double-mutant tetramer, where the two H88R chains are connected through a βC interface.

Trimers and dimers are denoted by a similar logic. Trimers are denoted as (3, *N*x, I). If *N* = 0 or 3, then I is not indicated. If *N* = 1, I denotes the relative position of the only H88R mutant chain with respect to the missing chain from the tetramer. If *N* = 2, I denotes the relative position of the only WT chain with respect to the missing chain. Furthermore, since we investigate dimer + monomer to trimer associations, the notion (I_dimer_)(3, *N*x, I_mut_) can also be applied here. Finally, dimers and dimer-formation reactions are denoted by (2, *N*x, I), where I denotes the monomer + monomer association interface (regardless of *N*).

### Model Building and Evaluation

Linear models were built to assign a Δ*G* contribution to structural components or effects inside the TTR tetramer. These models differ in the number of components and their role in tetramer formation. As usual, models with fewer parameters generalize and reflect component Δ*G* relationships well, but have a higher predictive error (bias-variance trade-off). Mathematically, these models correspond to matrices **M**, whose *M_ij_* elements describe the count-change of the *j*^th^component in the *i*^th^ association reaction. If **p** is the vector containing the model parameters (the component-wise Δ*G* values) and **g** is the vector containing the results of the MM/GBSA calculations (Δ*G* values corresponding to association-reactions), then we search for the best solution of the equation **Mp** = **g**, given a known **M**, **g** and a variance vector var(**g**). We chose to solve this equation using the weighted Moore-Penrose pseudoinverse approach^52^, i.e. to minimize the following weighted squared error:

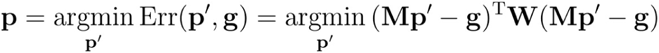

we calculate the following:

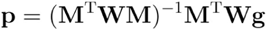

where **W** = δ*_ij_* / var(*g_i_*) is a diagonal weight matrix containing the reciprocal of the variances.

The simplest constructed model **M**^(1)^ counts the number of trivial interface formations in the association reactions. It only discriminates three components in the complexes, namely the βE, βC and lL interfaces, and associates a Δ*G* contribution to each of them. It disregards any mutations in the complexes, giving us only a general estimation for the interface Gibbs-free energies Δ*G*(βE), Δ*G*(βC) and Δ*G*(lL). As an example, during the (βE)(4, 2x, lL) association reaction 0 βE, 2 βC and 2 lL interfaces are formed.

In contrast to model **M**^(1)^, the second model **M**^(2)^ also takes mutations into account. There can be either 0, 1 or 2 mutations on a single interface, depending on the nature of the two chains forming that interface. This model differentiates 9 parameters; Δ*G*(βE0x), Δ*G*(βC0x), Δ*G*(lL0x), Δ*G*(βE1x), Δ*G*(βC1x), Δ*G*(lL1x), Δ*G*(βE2x), Δ*G*(βC2x) and Δ*G*(lL2x).

Finally, model **M**^(3)^ takes non-interface related parameters into account. As pointed out by Krissinel *et al*.^17^, entropic contributions often cannot be assigned to different components in protein complexes, but they are rather coupled to the nature of the whole complex (e.g. mass, symmetry, inertia). To model such interface independent energetic elements in the association Gibbs free energy, **M**^(3)^ considers every parameter of **M**^(2)^ with two additional parameters; Δ*G*(ch3), which corresponds to the trimer formation energy and Δ*G*(ch4), which corresponds to the tetramer formation energy.

### Protein Expression and Purification

Recombinant TTR variants were expressed and purified as described previously^35^. Briefly, the codon-optimized DNA sequence of mature TTR for the E. coli tRNA pool was synthesized (IDT) and cloned into the pGEX4T1 vector using BamHI and XhoI sites, generating an N-terminal GST-tagged construct with a thrombin cleavage site. Successful cloning was confirmed by Sanger sequencing. The plasmid was transformed into *E. coli* BL21(DE3) cells and used to inoculate LB pre-cultures (100 mg/L ampicillin), followed by 1 L TB fermentation at 37 °C, 180 rpm. Protein expression was induced at OD₆₀₀ = 0.8 with 0.1 µM IPTG and continued for 16 h at 28 °C. Cells were harvested, lysed by sonication in PBS containing 50 mM L-Arg/L-Glu (pH 7.3) and centrifugated. The supernatant was filtered and purified on a GSTrap affinity column; bound protein was eluted with 20 mM glutathione. The eluate was dialyzed, digested overnight with bovine thrombin (25 °C), and re-applied to the GSTrap column to remove the tag. The flow-through was concentrated and further purified by size-exclusion chromatography (Superdex 75, PBS pH 7.3, 1 mL/min). Fractions corresponding to the desired oligomeric state were pooled, verified by SDS-PAGE, aliquoted, flash-frozen in liquid nitrogen, and stored at -80 °C. The produced protein contains an extra glycine and serine on the N-terminal compared to the mature TTR protein (NCBI Reference Sequence: NP_000362.1)

### Peptide Mapping-based Tetramer Hybridization Experiment

To investigate the H88R/WT TTR hybridization, a mixture of H88R (monomeric) and WT TTR (tetrameric) was prepared containing 0.6 mg/mL of each variant in phosphate buffered saline (PBS) supplemented with sodium azide (50 μM) and dithiothreithol (DTT, 1 mM). Control experiments contain a single protein component with the same setup and experimental conditions. Then, the samples were incubated for one week at 37 °C. After incubation, samples were processed by size exclusion chromatography. Samples were concentrated down to 0.3 mL using Millipore Amicon 10 kDa MWCO ultracentrifugal filters (Merck, Germany) and injected to a Superdex 75 Increase 10/300 (Merck, Germany) column equilibrated with PBS (pH = 7.3). The flowrate was set to 0.75 mL/min, and fractions were manually collected based on UV absorbance measured at 280 nm. Calibration of the chromatographic system was done with a Gel Filtration Standard (Bio-Rad Laboratories, USA). Tetrameric fractions were further analyzed using peptide mapping. For peptide mapping, 10-10 μg of samples (based on absorbance measurements at 280 nm) were topped to 30 μL with PBS buffer. Protein samples for peptide mapping were denatured with RapiGest (Waters, Milford, MA, USA) at a final concentration of 0.05 %. Digestion took place with sidechain-protected porcine trypsin (Promega Corporation, Madison, WI, USA) at a 1:25 enzyme:protein ratio, at 37 °C and with shaking (300 rpm) overnight in an Eppendorf Thermomixer C thermoshaker. Sample cleanup was carried out on C18 spin columns (Pierce™ C18 Spin Columns, Thermo Fisher Scientific) as per the manufacturer’s instructions. The eluted samples were vacuum dried at 45 °C. LC-MS^E^ measurements were performed on a Waters Select Series IMS QToF Spectrometer coupled with an Acquity UPLC I-Class PLUS (Waters Corporation, Milford, MA, USA). Leucine-enkephalin was used as the lock mass for an accurate mass determination. Peptides were separated on a Waters Acquity UPLC Peptide CSH C18 column (1×150 mm, 1.7 μm). The autosampler and the column were maintained at 8 °C and 45 °C, respectively. Mobile phase A was water with 0.1 % (V/V) formic acid, and B was acetonitrile with 0.1 % (V/V) formic acid. Dried samples were dissolved in 20 µL of 5 % acetonitrile/water, 0.1 % (V/V) formic acid solvent mixture and 10 μL were injected. The following gradient program was used at a flow rate of 20 µL/min: 0–1 min, 5 % B; 1–45 min, 5–35 % B; 45–46 min, 35–85 % B; 46–50 min, 85 % B; 50-51 min, 85-5 % B; and 51-60 min, 5 % B. Source parameters were as follows: polarity, positive; capillary voltage, 2.7 kV; cone voltage, 40 V; source offset, 10 V; source temperature, 120 °C; desolvation temperature, 400 °C; cone gas flow, 20 L/h; desolvation gas flow, 800 L/h; and nebulizer gas pressure, 6 bar. The data acquisition was carried out in the 50–2000 m/z range in V-mode. Fragmentation was performed in the transfer cell, collision energy (CE) was ramped between 19 and 45 V. Data analysis has been performed in MassLynx 4.1 and BioPharmaLynx 1.3.5 (Waters Corporation, Milford, MA, USA).

### Patients

In this study we collected sera samples and clinical information of 3 patients carrying the mutation H88R with their informed consent in compliance with World Medical Association (WMA) Declaration of Helsinki. Ethics approval was received from National Centre for Public Health and Pharmacy (Hungary), reference number NNGYK/53227-6/2025. All examination and sample collection were carried out at Semmelweis University, Department of Internal Medicine and Hematology

Patient 1 (female) was diagnosed after two years of heart failure symptoms with ATTR-CM, NYHA stage II heart failure, bilateral CTS and paroxysmal atrial fibrillation and vitreous opacities at age 68. Her TTR serum level at the time of the diagnosis was measured 103 mg/L, before receiving any specific treatment. Her sera samples analyzed in this study were also collected before treatment. Later, she was enrolled in the ongoing Magnitude clinical trial (receiving CRISPR/Cas9 based TTR gene editing therapy vs. placebo, ClinicalTrials.gov Identifier: NCT06128629).

Patient 2 (female) and Patient 3 (female) are sisters aged 20 and 25. They are completely asymptomatic and free of complaints, with no significant medical history; their ECG and echocardiographic findings are normal. Their father was diagnosed at age 48 with H88R-mutated ATTRv, NYHA Class II heart failure, and Stage I Familial Amyloid Polyneuropathy (FAP). He received tafamidis treatment. Due to progressive heart failure, he underwent a heart transplant at age 52 and passed away during the early postoperative period. Their genetic screening was based on their family history and indicated that they are both H88R mutation carriers. After diagnosis, their sera were collected, and TTR level measured (Patient 2: 132 mg/L; Patient 3: 62 mg/L).

### Variant Ratio Analysis of Heterozygous H88R Carrier Patients’ Sera

LC-MS grade formic acid (FA), HPLC grade trifluoracetic acid (TFA), ammonium bicarbonate (≥99.5%), iodoacetamide (IAM) (≥99%) were purchased from Merck (Rahway, NJ, USA). Pierce™ top 12 abundant protein depletion spin columns and Pierce™ C18 spin columns were procured from Thermo Scientific (Waltham, MA, USA). LC-MS grade water, acetonitrile (ACN), and methanol were obtained from VWR (Radnor, PA, USA). DL-dithiothreitol (DTT) was purchased from Agilent Technologies (Santa Clara, CA, USA). RapiGest™ SF surfactant was acquired from Waters Corporation (Milford, MA, USA). Endoproteinase Glu-C was provided by Promega (Madison, WI, USA). PD MidiTrap™ G-25 size exclusion chromatography (SEC) columns were obtained from Cytiva (Marlborough, MA, USA).

For nano-liquid chromatography-tandem mass spectrometry (nano-LC-MS/MS) analyses, blood serum samples were processed as follows. Abundant serum proteins were depleted using Pierce™ top 12 abundant protein depletion spin columns based on the manufacturer’s protocol. Briefly, 10 µL of serum was added to the column equilibrated to 25 °C, then incubated in a thermoshaker for 60 min at 25 °C while continuously shaking at 900 rpm to ensure that the sample mixes with the slurry. The bottom closure and the cap were removed from the column, and the filtrate was collected into a LoBind Eppendorf tube by centrifugation (3,000 × g, 2 min, 25 °C). The volume of each sample was adjusted to 1 mL, then solvent exchange was performed on PD MidiTrap™ G-25 SEC columns. The columns were equilibrated with 20 mL of LC-MS grade water, and the samples were added to the column. After the liquid entered the resin completely, 1.5 mL water was added, and the macromolecular (protein) fraction was collected into a low-binding tube. The volume of each sample was reduced to 100 µL in a vacuum concentrator at 60 °C. Thereafter, 4.5 µL (0.5%) of RapiGest™ SF surfactant and 1.5 µL (200 mM) of DTT were added to the samples and incubated at 95 °C for 30 min. After cooling down to room temperature, 13.1 µL (200 mM) of ammonium bicarbonate and 3 µL of iodoacetamide (IAM) were added, followed by incubation at room temperature for 30 min in the darkness. The alkylation was quenched by adding 1.5 µL (200 mM) DTT. The samples were then digested by adding 1.2 µL (1 µg/µL) of endoproteinase Glu-C and incubated for 16 h at 37 °C with gentle mixing at 250 rpm. The digestion was quenched by adding 3 µL of FA and incubating for 30 min at 37 °C. The samples were then dried in a vacuum concentrator at 60 °C, reconstituted in 100 µL of 0.1 % TFA, and desalted by means of Pierce™ C18 spin columns. In all steps, the flow of liquids through the column was ensured by centrifugation for 2 min at 1800 × g. The columns were activated by adding 200 µL of 50% methanol twice, washed with 200 µL of 5% ACN containing 0.5% TFA twice, and equilibrated by adding 0.1% TFA twice. 50 µL of the sample was applied to the column, and the flow-through collected into a new low-binding tube was reloaded after centrifugation. The column was then washed with 100 µL of 0.1% TFA twice, then the peptides were eluted twice into a new collection tube with 50 µL of 70 % ACN containing 0.1 % TFA. The eluate was then dried in vacuum concentrator at 60 °C and kept at −20 °C until further analysis.

The nano-LC-MS/MS measurements were performed on a Waters nanoAcquity ultraperformance liquid chromatography (UPLC®) system (Milford, MA, USA) coupled to a Bruker Maxis II ETD Q-TOF (Bremen, Germany) MS. The column thermostat and the autosampler were maintained at 50 °C and 14 °C, respectively. The peptides were trapped on a Waters Acquity UPLC® M-Class Symmetry® C18 trap column (180 µm × 20 mm, 100 Å, 5 µm) and separated on a Waters Acquity UPLC® M-Class Peptide BEH C18 analytical column (75 µm × 250 mm, 130 Å, 1.7 µm). Mobile phase A was 0.1 % FA in water, while B was 0.1 % FA in ACN, respectively. For the measurements, dried samples were reconstituted by sonication for 3 min in 20 µL of 2 % ACN containing 0.1 % FA. The resulting solution was transferred into autosampler vials, from which 1 µL was injected into the chromatography system. Trapping was performed with 99 % A at a flow rate of 10 µL/min for 3 min. For peptide separation, the gradient program at a flow rate of 0.3 µL/min was as follows: 0-1 min, 12 % B; 1-34 min, 25 % B; 34-54 min, 45 % B; 54-64 min, 95 % B; 64-74 min, 95 % B; 74-74.5 min, 1 % B; 74.5-100 min, 1 % B. Before every analytical run, 5 µL of sodium formate calibrant (15 mM) was injected into the chromatography system.

Sample ionization was achieved via a CaptiveSpray electrospray (ESI) ion source equipped with nanoBooster, operating in positive mode. The source parameters were as follows: capillary voltage, 1150 V; nanoBooster pressure, 0.35 Bar; drying gas flow rate, 3 L/min; drying gas temperature, 160 °C. Ion transfer parameters were as follows: funnel 1 RF, 400 Vpp; multipole RF, 800 Vpp; collision RF, 1,200 Vpp; transfer time, 120 µs; pre pulse storage, 10 µs. MS/MS data was obtained using data dependent acquisition (DDA), with a fixed cycle time of 2.5 sec. MS spectra were acquired in the range of 150-2200 *m/z* at 3 Hz. Collision-induced dissociation (CID) was performed at 16 Hz for abundant precursors, and at 4 Hz for low-abundance ones. Collision energy (CE) was determined by the control software based on the *m/z* value of the precursor ions. Following selection for fragmentation, precursors were excluded from further fragmentation for 2 minutes, unless their intensity increased threefold. For mass calibration of precursor and fragment ions, data was recalibrated by the Compass DataAnalysis software 4.3 (Bruker Daltonics, Bremen, Germany).

Compass DataAnalysis and Byonic v5.0.20 (Protein Metrics, Cupertino, CA, USA) were used for data analysis and database search. A FASTA file of human transthyretin (UniProt entry: P02766, accession date: 2025-05-30) was generated and then modified to include a version with histidine to arginine substitution at position 108 (residue 88 in the mature peptide sequence), corresponding to the H88R variant. This FASTA file was used for the database search with Byonic using the following parameters: cleavage site, E; cleavage side, C-terminal; digestion specificity, fully specific; missed cleavages, 2; precursor mass tolerance, 10 ppm; fragment mass tolerance, 20 ppm; fragmentation type, QTOF/HCD; carbamidomethyl as fixed modification; oxidation, deamidation on N, deamidation on Q, and acetylation as variable modifications.

## Results and Discussion

### In vitro Identification of H88R Hybrid Tetramers

Size exclusion chromatography (**Figure 2**, **Panel A**) of the TTR H88R sample after the incubation period revealed aggregation and possible fragmentation that led to the complete disappearance of the monomeric peak. In contrast the chromatogram of TTR WT showed no aggregation and was completely tetrameric. The mixture of TTR WT and H88R, however, showed some aggregation less than that of the pure TTR H88R, an increase in the area of the tetrameric peak relative to both the freshly made mixed control sample and the incubated pure TTR WT sample, and a small peak of monomers. Taking these observations altogether, we argue that TTR H88R does form hybrid tetramers with the WT variant, which in turn helps to prevent aggregation and possible fragmentation events, leading to lower aggregation rate and an observable peak of monomeric TTR H88R. Furthermore, tetrameric peaks of the WT sample and the freshly prepared H88R+WT mixture sample were smaller than the tetrameric peak of incubated H88R+WT mixture, strongly suggesting the incorporation of H88R TTR into the tetramers. Peptide mapping of this tetrameric peak using mass spectrometry strongly proved the presence of H88R TTR (**Figure 2**, **Panel B**).

**Figure 2.**
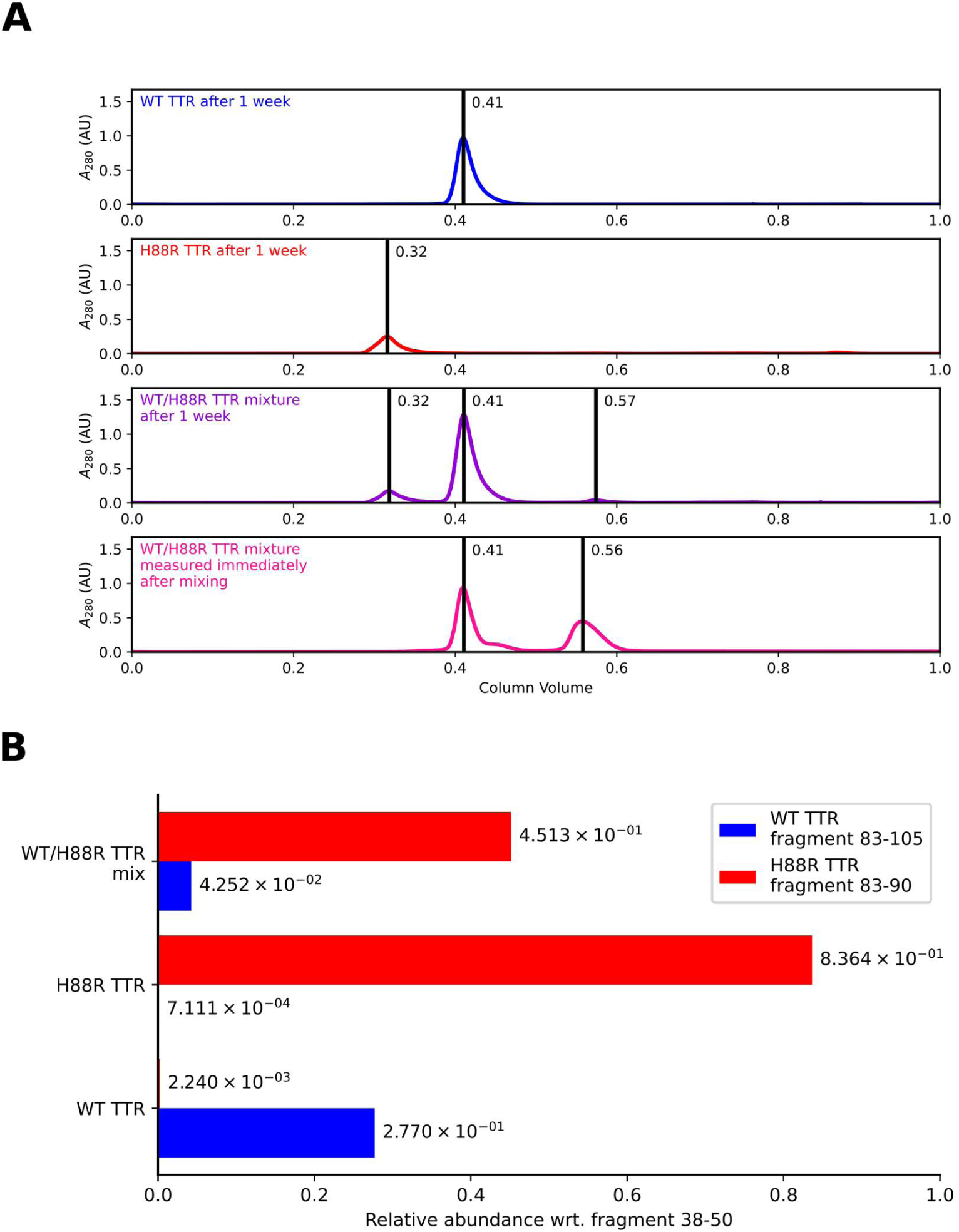
Results of the hybridization experiment. Panel **A** shows the size exclusion chromatograms of the samples. Peaks at 0.32 CV (Column Volume) correspond to aggregates, 0.41 CV to tetramers and 0.56 to monomers. Tetrameric peaks were collected and analyzed by tryptic peptide mapping on an LC-MS instrument. Panel **B** depicts the relative abundance of tryptic peptides representative of the WT and H88R TTR chains compared to the most intensive peptide contained in both variants (38-50). Numbering in this instance starts with the two leftover residues of the N-terminal thrombin cleavage site (Gly-Ser). For this reason, residue numbers are increased by two compared to the mature peptide sequence.

### Secretion of H88R TTR is Hindered in Patients

In the nano-LC-MS/MS workflow, abundant proteins were first depleted from the serum samples. After solvent exchange, the proteins were reduced, alkylated, digested with endoproteinase Glu-C, and desalted prior to the analytical runs. Glu-C was selected for digestion to generate peptides similar in size around the amino acid substitution facilitating comparison of the variants. Database searching with Byonic confirmed the presence of both variants with high confidence, yielding Byonic scores of 574 and 454 and LogProb values of 7.05 and 5.77 for the WT and H88R variants, respectively, for the 16-amino-acid peptides E.IDTKSYWKALGISPFHE.H (**Supplementary Figure 4**) and E.IDTKSYWKALGISPFRE.H (**Supplementary Figure 5**) detected in the spiked sample (**Supplementary Figure 6**). As expected, the peptide derived from the H88R variant exhibited a slightly lower retention time in reversed-phase chromatography than the peptide derived from the WT protein. Manual inspection of the extracted ion chromatograms identified these peak pairs in the studied samples based on retention behavior and mass accuracy (0.00–2.98 ppm), indicating the presence of the H88R variant. (**Figure 3**) In all samples, peak area of H88R derived peptides was much smaller (12.5-16.5 times smaller) than that of the WT derived peptide. This proves what we proposed earlier based on observed commonalities with TTR variants D18G and A25T that secretion of H88R TTR into the serum is greatly reduced^35^.

**Figure 3.**
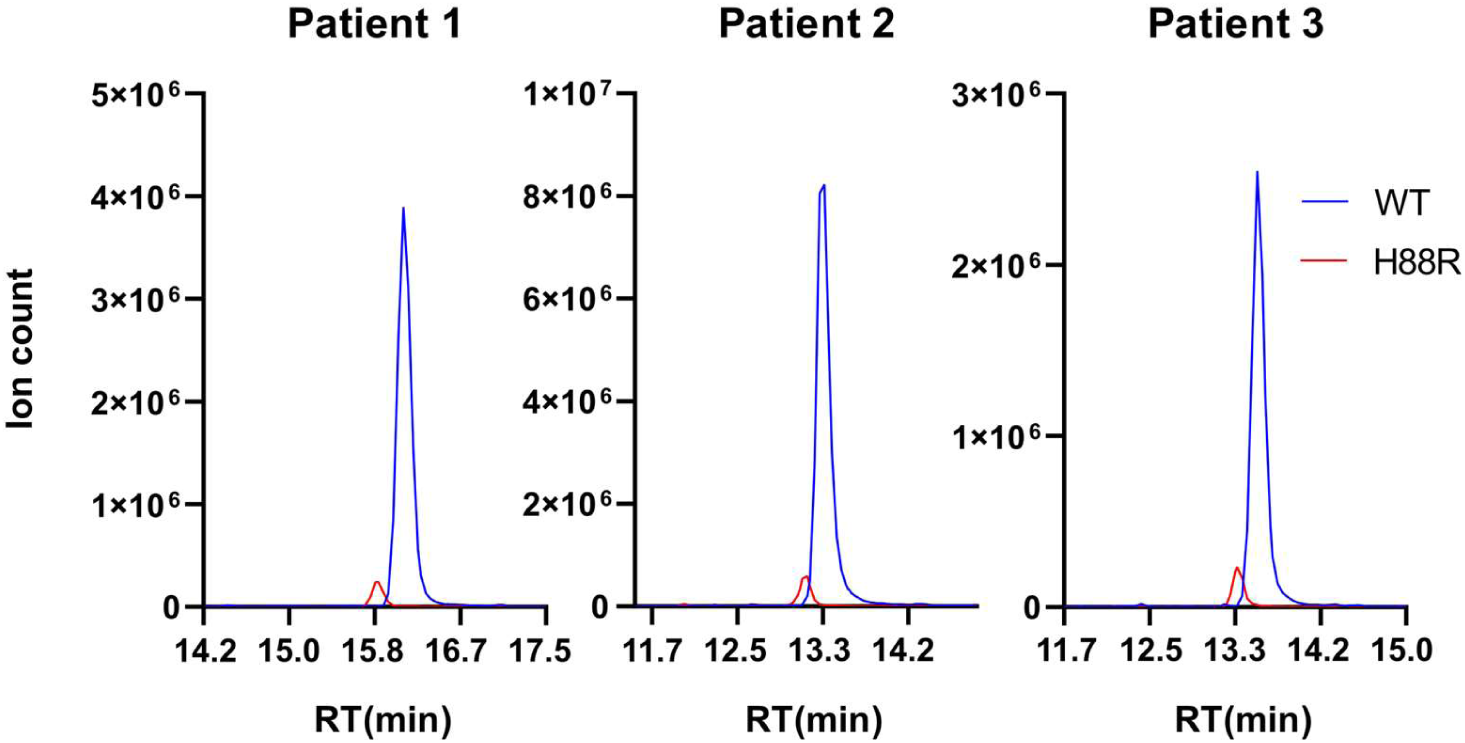
Extracted ion chromatograms of digested serum LC-MS/MS measurements. Red chromatograms are showing the elution of E.IDTKSYWKALGISPFRE.H peptide representing the H88R variant content of the samples, while blue chromatograms show the elution of the E.IDTKSYWKALGISPFHE.H peptide representing the WT variant TTR content of the samples. All three patients exhibited strongly decreased H88R serum content similar to patients with D18G and A25T genotype described by others.

### Modeling the Interaction of H88R TTR Monomer with the Endoplasmic-Reticulum-Associated Protein Degradation Machinery

To grasp a better understanding of the fate of the H88R TTR monomer within our physiology, we investigated whether it might form an association with the BiP chaperon of the Endoplasmic-Reticulum-Associated protein Degradation (ERAD) pathway leading to its retention in the ER and subsequent proteasomal degradation, as was demonstrated in case other monomerizing variants, such as D18G and A25T TTR^53–55^.

Recently, the structure of the nucleotide binding domain of BiP (residues 25-405) in complex with cerebral dopamine neurotrophic factor (CDNF) has been determined (PDB ID: 9H0C^56^). The CDNF segment that associates with the ATPase (nucleotide binding) domain of BiP is composed of a Schellman helix-C-capping motif, very similar to the EF-helix/EF-loop region of TTR, which we previously identified as a key organizational center of its monomeric units^35^. In place of W79 of TTR (the central residue of the motif), CDNF carries a tyrosine (Y^141^). The EF-helix/EF-loop region (residues 75-87) also partially overlaps with the tryptic fragment of D18G TTR (residues 81-103) that was confirmed to bind to BiP using MS/MS measurements^57^.

The overall structural similarity of the two segments of CDNF and TTR allows a prediction of a similar binding conformation for the latter too. The notion that monomeric TTR variants might bind to the same site of BiP as CDNF is best supported by the finding that incubation of the BiP / D18G TTR complex with ATP leads its complete dissociation^58^. The ADP/ATP nucleotide exchange induces significant structural rearrangement in BiP (PDB ID: 6ASY^59^): the previously unstructured linker segment (residues 414-421) becomes ordered and forms a β-strand that fills the same binding pocket that CDNF was found anchored to (and where we propose the monomeric TTR variants might also bind), while the β-barrel of the substrate binding domain (residues 420-500) completely blocks its entrance. Although in this specific BiP/CDNF complex BiP contains only a phosphate group, glycerol, Mg^2+^ and Cl^-^ ions at the nucleotide binding site, it is nearly isomorphic with the ADP-bound, ligand-coordinating conformation of the chaperone (PDB ID: 3IUC^60^). Using these two crystal structures, it was thus possible to create a BiP-TTR complex structure.

This proposed binding mode does not allow for the chaperone to coordinate the tetrameric form of TTR (which would cause severe clashes), and we also found that it is able to distinguish in favor of the H88R variant. MCMM calculations were used to optimize the binding conformation of WT and H88R TTR monomers, the latter carrying the open conformation of the EF-helix/EF-loop segment that it also samples (according to our previous results^35^). We found that this open conformation was able to form a much closer fit with the binding pocket of BiP, resulting in 4 additional H-bonds between the two (see **Figure 4**). The H88R variant chain also forms (in contrast to the WT chain) an additional contact with BiP/R^197^, a residue which was also found to be a key interaction regulator between BiP and other chaperons^61^ and co-chaperons^62^. Accordingly, the binding energy of the complexes (estimated by single-point MM/GBSA) were found to be -8.03 kcal/mol and -19.65 kcal/mol, for the WT and H88R variants, demonstrating that BiP might indeed be able to select the mutant and assist in its retention within the ER. It is interesting to note that neither the previously considered D18G and A25T mutations, nor the presently investigated H88R mutation site is part of the actual BiP binding region of TTR, they simply increase the pliability and accessibility of the implicated residues by destabilizing the tight fold of the monomeric structure and especially of the EF-helix/EF-loop region^35^. Hammerström et al. have reported that the tryptic fragment containing residues 81-103 was able to partially displace flag-tagged D18G TTR from its complex with BiP^63^, showing that removal of the structural restraints exerted by the fold enhances the binding affinity of the unstructured tryptic fragment, in nice agreement with our finding that the H88R TTR monomer with its open and accessible EF-helix/EF-loop region forms a much stronger complex with the chaperone than the WT variant.

**Figure 4.**
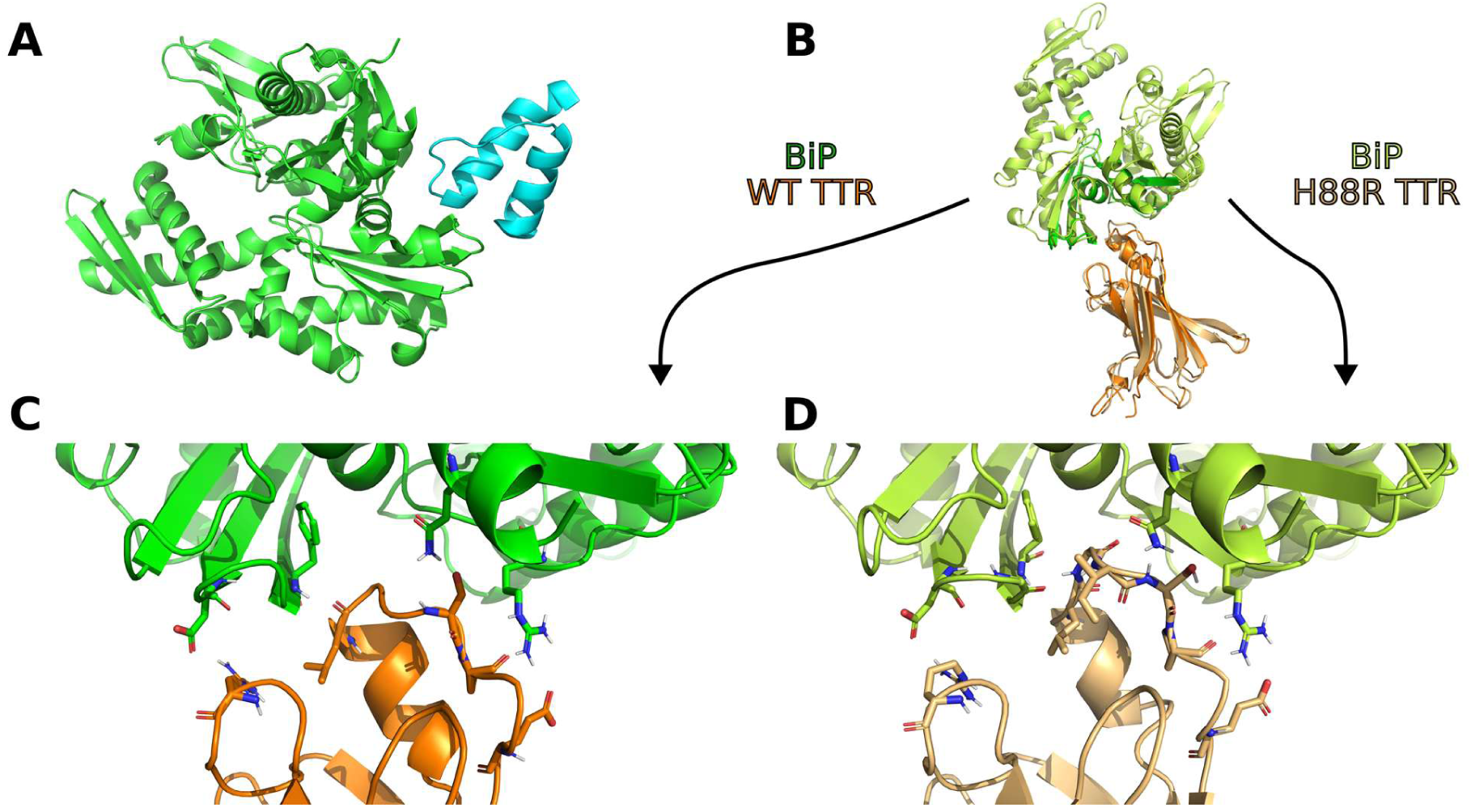
On **Panel A**, the structure of the recently published CDNF/BiP complex can be seen (PDB ID: 9H0C, BiP is shown in green, CDNF is shown in cyan). On **Panels B**, **C** and **D** the MCMM results are visible for the proposed TTR/BiP complexes. **Panel B** shows the WT and H88R variants overlaid with each other, while **Panels C** and **D** shows close-up images of the binding regions of the WT and H88R mutant cases, respectively. TTR is colored in orange (WT) and in light-orange (H88R). Contacting residues are highlighted as sticks. Beside the TTR/R^21^-BiP/D^238^, TTR/L^82^-BiP/F^242^, TTR/S^85^-BiP/Q^401^, TTR/P^86^-BiP/R^197^ and TTR/E^89^-BiP/R^197^ H-bonds of the WT-TTR/BiP complex, in case of the H88R variant of TTR two further backbone-backbone H-bonds (TTR/G^83^-BiP/Q^240^ and TTR/I^84^-BiP/Q^240^) and two further connections between TTR/(P^86^, E^89^) and BiP/R^197^ are formed.

### Assessing the Stability of Hybrid (WT/H88R) Tetramers: Gibbs Free Energies of Association Reactions

To map the possibilities of hybrid complex formation between the WT and H88R monomer of TTR, MD simulations of all considered assemblies were carried out, followed by MM/GBSA analysis of the trajectories. **Supplementary Figure 7** shows the MM/GBSA predicted Δ*G* value of each association reaction. It can be seen that TTR monomer dimerization-, monomer-dimer association-, and dimer dimerization reactions were considered. Furthermore, different hybrid formation patterns were also investigated. Assuming two different chains (WT and H88R) present in the solution, three different interaction modes (βE, βC, lL) between chains, and equivalence of chains in the complexes, one can construct nine different TTR dimers, eight different trimers and seven different tetramers. As can be seen from the MM/GBSA data, generally, the largest amount of energy is freed when two βC dimers form a tetramer, while, logically, the smallest amount of energy is freed during monomer dimerization reactions. To draw conclusions about the role of different structural components present in these oligomers, we constructed three models (**M**^(1)^, **M**^(2)^, and **M**^(3)^) to extract energetic contributions that build up the full association Δ*G* values. The appropriate interpretation of these energetic contributions allows us to suggest pathways along which hybrid TTR oligomers can be built up, and to make assumptions about the equilibrium ratio of different hybrid oligomeric TTR species.

In model **M**^(1)^, which only discriminates the βE, βC and lL interfaces, we can extract the general energetic content of these three different TTR contact surfaces. It is important to keep in mind, however, that these energies do not belong to specific complexes and only indicate tendencies. In other words, these are the marginalized expectations of interface energies over every possible complex and mutation state. Results and fitting statistics for **M**^(1)^ are summarized in **Table 1**. Note that χ^2^ >> 1, indicating that the model is yet highly underfitted, possibly due to the low number of parameters. It is clearly visible that the strongest interface in these complexes is the βE interface, followed by the lL and βC interfaces. The difference behind their assigned Δ*G* values is significant. These results also reflect intuitive expectations, since the βE interface is by far the most extended. Several strong backbone-backbone H-bridges hold its two anti-parallel oriented β-strands in close proximity, in addition to hydrophobic side chain interactions. The βC interface contains the large ligand binding cavities of TTR. To allow the binding of ligands in between them, the βC surfaces are loosely packed, thus it is unsurprising that this interaction was shown to be the weakest in our calculations. In the ligand-free state these hydrophobic cavities are filled with water molecules, creating unfavorable apolar-polar interactions between the protein and the solvent. Possibly, this contributed to the low energy gain when a βC interface is created.

**Table 1.**
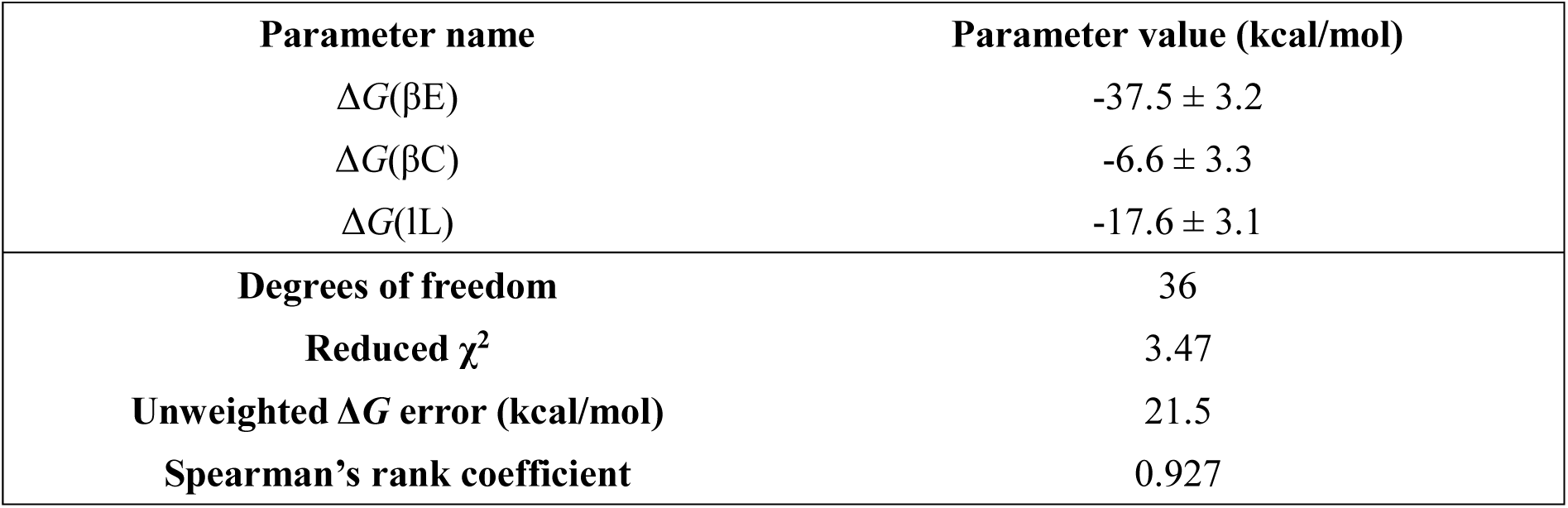
Table showing the **M**^(1)^ model parameters and fitting statistics, where interface mutation states are ignored. A general tendency for the interface binding strengths can be observed; the formation of a βE interface is the most favorable, followed by the lL and βC interfaces. Δ*G* differences are significant. The high χ^2^ score indicates underfitting.

Besides the different interface types, model **M**^(2)^ also takes mutation information into account. The nine different parameters split every dimer-dimer interface type into WT vs. WT (0x), WT vs. mutant (1x), and mutant vs. mutant (2x) chain-chain contacts. In other words, we can investigate the effects of mutations on different interfaces, generalized to all (dimer, trimer and tetramer) complexes. Results and fitting statistics for **M**^(2)^ are summarized in **Table 2**. The same tendency as in **M**^(1)^ for the interfaces is visible, which is a good retrospective validation of the built model. Although we employed more parameters in **M**^(2)^ than in **M**^(1)^, the unweighted fitting error and χ^2^ became slightly worse. The only significant effect observable from these data is the change between Δ*G*(βE1x) and Δ*G*(βE2x). It can be seen that two H88R mutations on the same βE interface highly decrease the energy gain during complex formation. As later can be seen, this is the result of a complex steric, electrostatic and allosteric interplay between the R88 and its environment.

**Table 2.**
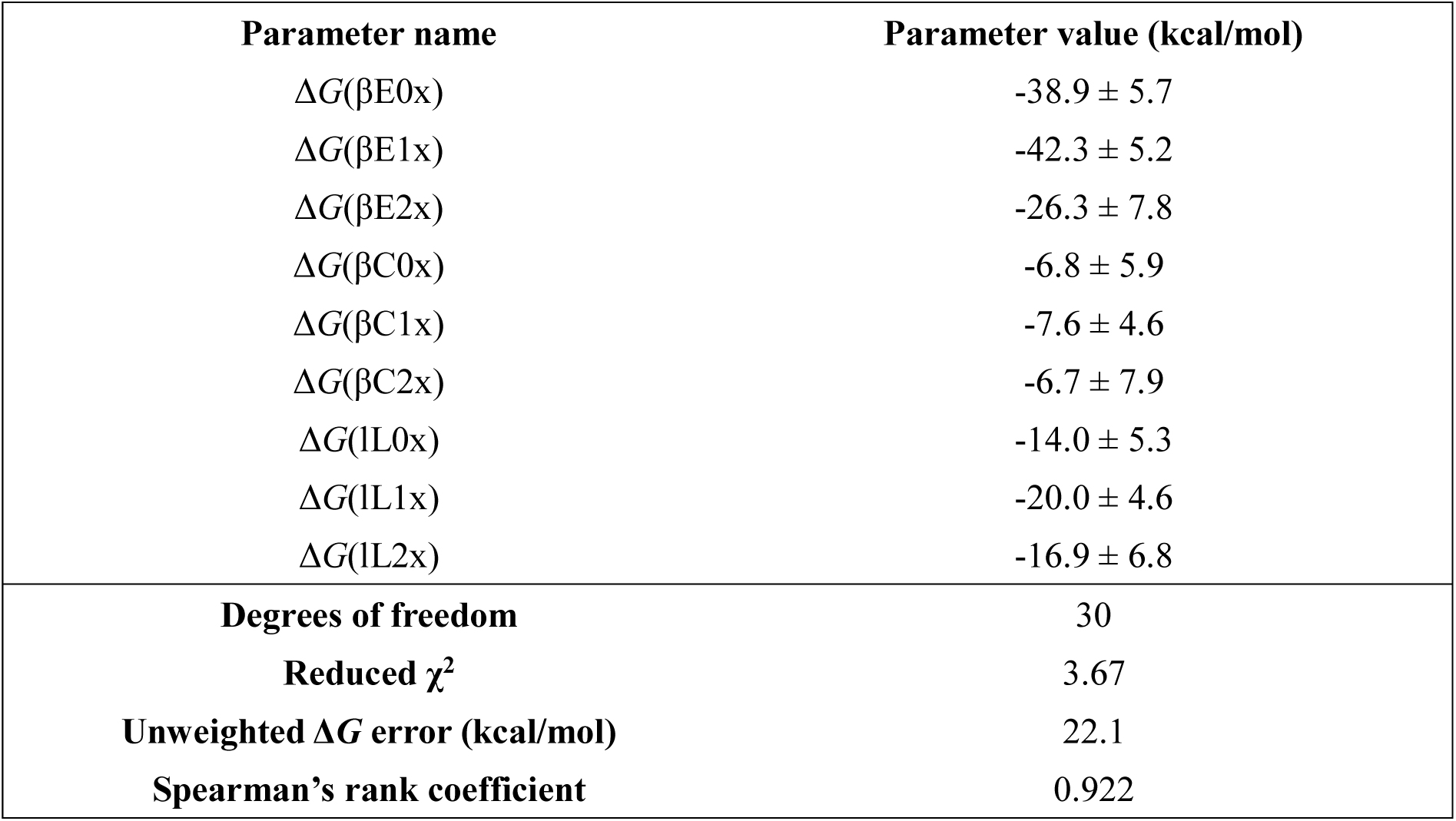
Table showing the **M**^(2)^ model parameters and fitting statistics, where interface mutation states are already taken into account. The previous tendency for the interface binding strengths can still be observed, but relations between the same interfaces with different mutation states can also be examined. The only two significant, same-interface Δ*G* difference belongs to Δ*G*(βE2x). Two H88R mutations on the same βE interface seem to inflict disadvantageous interactions. According to the χ^2^ score, the model still underfits, even though the number of parameters tripled.

**M**^(3)^ is a rational correction to **M**^(2)^, which takes complex-specific effects into account that could not be explained with chain-chain interaction components. This model contains interaction Δ*G* values, as well as specific correction factors for tetramers (Δ*G*(ch4)) and trimers (Δ*G*(ch3)). During association reaction Δ*G* calculation, these shift the baseline energies of tetramers and trimers with respect to dimer energies (i.e. Δ*G*(ch2) is taken to be 0 kcal/mol, see **Table 3**). We can see a large reduction in the reduced χ^2^ value and the unweighted reaction Δ*G* error when compared with those of the previous models. Even though **M**^(2)^ triples the number of parameters of **M**^(1)^, we do not see as much of an improvement in these descriptors, as when we move from **M**^(2)^ to **M**^(3)^. This highlights the importance of complex specific energetic components. As expected, the energetic correction of trimers is less favorable than that of the tetramers. Interface-specific energies are also shifted, although their order remains the same. The βE interface energy is still highly favorable. The lL interaction only slightly and not significantly drives the association, while the presence of a βC interaction is predicted to push complexes apart. This is a more intuitive scale of values, as the lL interface is small and thus should have only a small effect on complex formation, while the βC interface, in its empty form, is a hydrophobic cavity filled with water, the formation of which should be discouraged. **Supplementary Figure 8** shows the correlation scatter plots between the MM/GBSA predicted Δ*G* values and the predictions of all three models. It can be seen that as model complexity increases, the correlation gets higher.

**Table 3.**
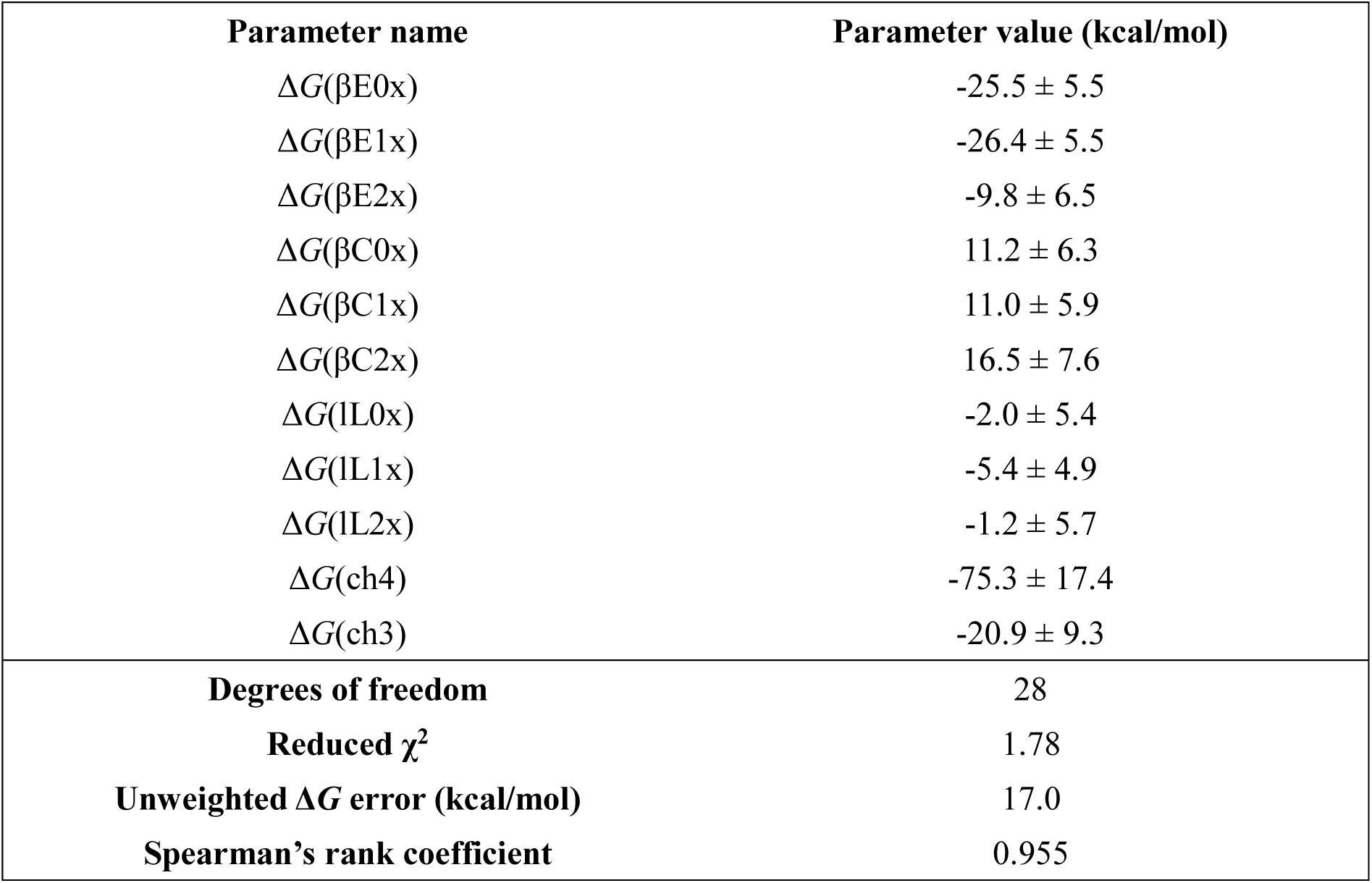
Table showing the **M**^(3)^ model parameters and fitting statistics, where both interface mutation states and complex-specific energy corrections are considered. Although only two additional parameters are introduced to **M**^(2)^ in order to obtain **M**^(3)^, the fitting error and χ^2^ score dropped by a high amount relative to the drop of **M**^(1)^ and **M**^(2)^. This indicates that complex-specific model parameters are necessary for the accurate description of these reactions.

### Structural Effects of the H88R Mutation in Tetramers

To investigate the structural effects of the H88R mutations on tetramer formation and connect them with the calculated binding energies, we also analyzed residue – residue interactions on different interfaces. For this, we utilized the Residue Interaction Network Generator (RING) software^49^, as it has proven to be useful for the analysis of protein complexes in our previous studies^50,64^. Both interaction statistics, as well as interaction graph visualizations were extracted from the TTR tetramer trajectories. Two types of interaction statistics have been included; first, the frame-average of different secondary chemical interaction counts, and second, the residue importance values based on the frame-average of secondary chemical interaction counts formed by that residue. Both statistics types were calculated for all three interface types. Results are collected in **Supplementary Tables 1** and **2**.

It can be seen that the substitution of H^88^ with R^88^ causes large changes both in interaction quantities and qualities on all interfaces. Considering the H-bonds on the βE interface, the WT and double mutant hybrid tetramers form the most of them (around 33-34, on average), the triple mutant forms H-bonds somewhat less frequently, and the single mutant and fully mutant homotetramers form the least amounts. In a similar way, the WT complex has the most van der Waals (VdW) contacts and π-π interactions, hybrid mutants display intermediate amounts, and the fully mutant complex has the lowest number of these interactions. Interestingly, ionic interactions (i.e. salt bridges) show different tendencies; while complexes (4, 0x), (4, 1x), (4, 2x, βE) and (4, 2x, βC) do not appear to form salt bridges, complexes (4, 2x, lL), (4, 3x) and (4, 4x) introduce either one or two transient ionic interactions.

According to the interaction counts, residues with the most contacts on the βE interface are F^87^, F^95^, Y^116^, and T^118^. In the WT tetramer, F^87^ is responsible for a multitude of inter-chain VdW contacts with F^95^, Y^105^, I^107^ and A^120^, in addition to a weak, but persistent backbone-backbone H-bond with T^96^ (**Figure 5**, **Panels A** and **C**). More importantly, although not interacting directly with it according to RING, its side chain is also close to the H^88^ side chain (the distance of A/F^87^ Cδ and A/H^88^ Nε is 3.8 Å in the medoid structure of the most populated cluster of the WT tetramer). Residue F^95^ is part of the same hydrophobic cluster as F^87^. In fact, close and interacting aromatic side chains pave the whole βE interface from end to end. At the core of the βE interface a pair of Y^116^ and a pair of T^118^ residues can be found, whose backbone-backbone H-bonds solidify the interaction between the two β-edges (**Figure 5**, **Panels B** and **D**). Additionally, in the WT structure T^118^ forms important side chain-side chain H-bonds with H^88^ in three out of four chains (the interaction of A/H^88^ and B/T^118^ is weak). In contrast, the average number of contacts significantly decreases with respect to the WT complex after the introduction of all four H88R mutations. In the WT tetramer four residues have 4.0 contacts or above, while in the fully mutant tetramer only has one residue with this property (D/F^87^). Residue A/F^87^ in complex (4, 4x) only interacts with F^95^, T^96^ and A^120^. The H-bond with T^96^ is retained, albeit in the mutant case sometimes a backbone-side chain Oγ1 interaction is preferred, instead of a backbone-backbone one.

**Figure 5.**
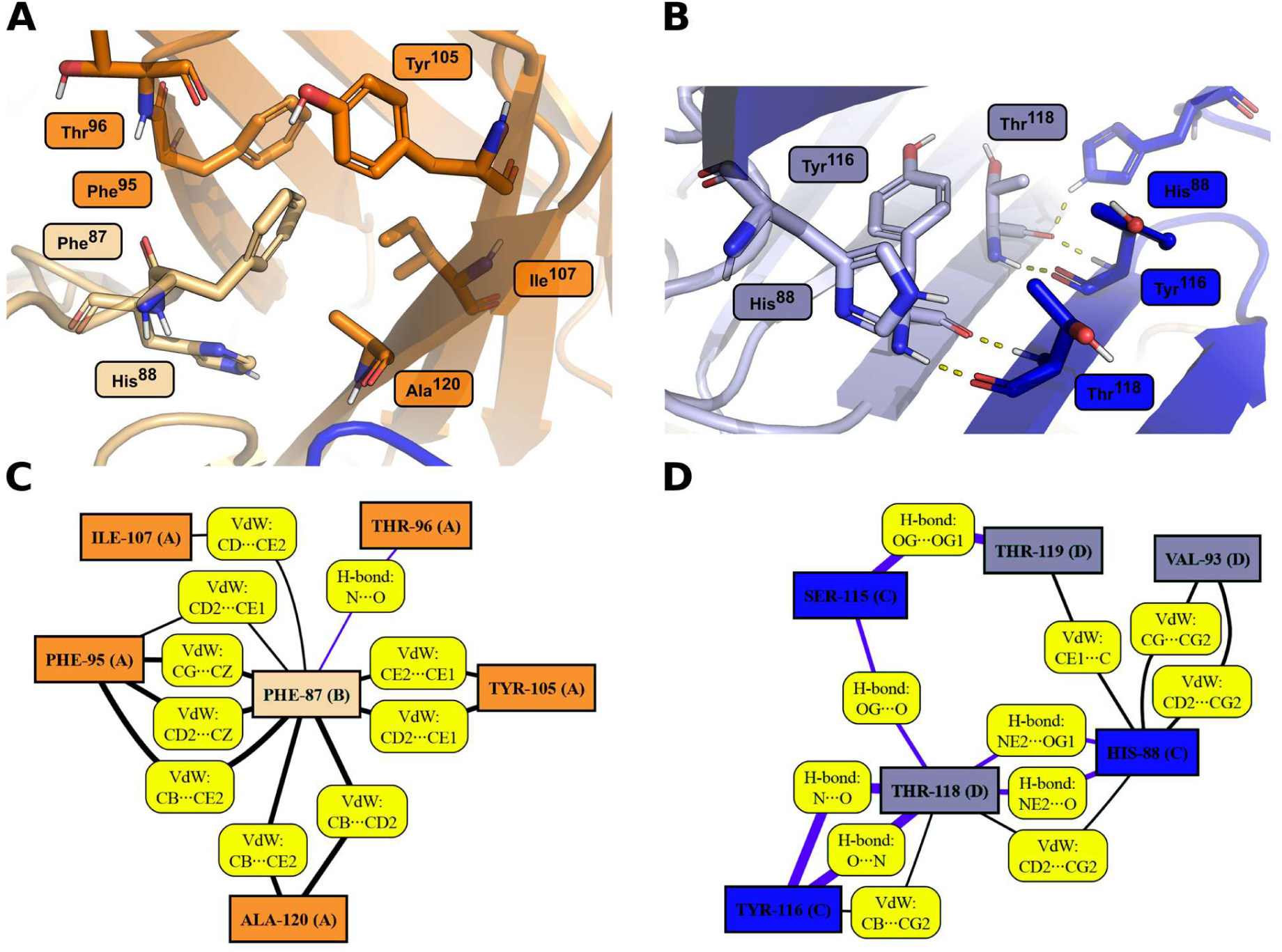
Panels **A** and **B** show important interactions in the WT TTR tetramer (i.e. complex (4, 0x)) in the first cluster center generated from the MD simulation trajectory. On panel **A**, the residues surrounding B/F^87^ can be seen, all of which form a hydrophobic pocket, important in the inter-chain interaction. On panel **B**, the core of the βE interface is highlighted, where three-three residues, namely H^88^, Y^116^ and T^118^ strengthen the inter-chain contacts. Besides the H-bonds between the backbone atoms, the H^88^ side chain can also hold onto the carbonyl or hydroxyl group of T^118^. Panels **C** and **D** show these interactions as graphs. Edge widths represent the strength of an interaction; wider edges mean stronger connection presence during the simulation. Blue edges are H-bridges, while black edges indicate VdW connections.

The direct effects of the histidine-arginine change on the βE interface can be observed from the average contact counts of the residues at position 88 (**Supplementary Table 3**). These effects show no clear tendencies; both H^88^ and R^88^ are capable of producing both robust and weak contacts, although the full average contact count for H^88^ (1.182) is slightly and not significantly lower than that of R^88^ (1.408), possibly due to the larger arginine surface area and its more numerous contact mode possibilities. In the TTR tetramers, H^88^ makes the most contacts in (4, 0x) chain C and reaches the least amount in (4, 3x) chain D. In the former case, C/H^88^ forms VdW contacts with V^93^, T^118^, T^119^ and A^120^. Additionally, a strong Nε2-Oγ1 or Nε2-O H-bridge is also present with T^118^ (**Supplementary Figure 9**, **Panels A** and **B**). In the latter case, D/H^88^ forms intramolecular H-bridges with S^112^ and S^115^ and faces away from the close C-chain residues V^93^ and F^95^ (**Supplementary Figure 9**, **Panel C**). R^88^ can be found in several different states in the complexes, which is visible from the χ_1_ side chain dihedral angle distributions (defined by the dihedral angle between atoms N-Cα-Cβ-Cγ, see **Supplementary Figures 10-16**). Different R^88^ side chain orientations populate the (4, 4x) complex; on chain A, the R^88^ side chain atoms arrange themselves approximately parallel to the chain A – chain B βE interface (χ_1_ ≈ 60° – 70°). On chain B, R^88^ has more freedom to move, as it escapes from the dense interface region and faces more towards the solvent (χ_1_ ≈ -180°). Residue C/R^88^ is also unique in this complex, since it is firmly held in place by intra-chain residues; it interacts with the backbone carbonyl of S^112^, P^113^ and S^115^, and also forms a π-cation interaction with Y^116^, whilst not making any contact with chain D (χ_1_ ≈ -60° – -70°). Finally, D/R^88^ orients itself similarly as A/R^88^ does, but with a highly different χ_1_ angle distribution (χ_1_ is bimodal with one, smaller mode around -60° – -70°, and with a more populated mode around -180°). These conformations are depicted on **Supplementary Figure 17**.

On the βC interface the largest count differences of H-bonds and VdW interactions are 1.0 (complexes (4, 0x) and (4, 3x)) and 4.1 (also for complexes (4, 0x) and (4, 3x)), respectively. This shows that the incorporation of multiple H88R mutations into the complex creates more contacts on the βC interface, which contrasts with the effects of these mutations on βE interfaces. However, this increase in contacts does not cause a significant decrease in calculated interface Gibbs free energies seen in models **M**^(2)^ and **M**^(3)^ (although it should be noted that these models also take trimer and dimer dissociation complex energies into account). Residues with the most contacts on the βC interface generally for all hybrid and non-hybrid complexes are L^17^, G^22^, T^119^, V^121^ and V^122^ (**Supplementary Figure 18**). In the WT complex, V^121^ mediates a ternary contact between three chains; A/V^121^ is connected to C/L^17^, C/G^22^, C/S^23^ and C/P^24^, and although it does not interact directly with them, it is also close to B/F^87^ and B/Y^114^. V^121^ is one of the “gate” residues at the ligand binding site of the βC interface and contributes to its hydrophobicity. Besides C/V^121^, A/G^22^ is participating in a strong backbone-backbone H-bond with C/V^122^ and, similarly to the case of V^121^, it is also part of the ternary βC-lL (i.e. A+C+D chain) interface (is close to B/Y^114^). L^17^ serves a highly similar purpose as V^121^ does, since it is also situated at the gate of the ligand binding site, although deeper.

On the lL interface the maximum count differences are 1.8 for H-bonds (complexes (4, 2x, βC) and (4, 3x)) and 3.6 for VdW contacts (complexes (4, 2x, lL) and (4, 3x)). It is interesting to see that while the complex (4, 3x) has the highest number of contacts on the βC interface, it also has the lowest number of contacts on the lL interface, indicating opposing mutation effects on these interaction surfaces. Residues Y^114^, A^19^, S^112^ and V^20^ form the majority of interactions on this interface. Y^114^ and A^19^ participate in both VdW contacts and backbone-backbone H-bridges. S^112^ forms inter-chain H-bonds with S^112^ of the neighboring chain and VdW contacts with A^19^. V^20^ also interacts symmetrically with another V^20^ residue and forms strong dispersive interactions with Y^114^.

### Dynamic Behavior of the Complexes

We used PCA- and RBS-based analysis methods to extract correlative motion information of all possible hybrid and non-hybrid TTR tetramers. Based on the MD trajectory, PCA is able to assign representative, concerted motion modes to the selected atoms. Principal components are ordered according to their amplitudes, where the first component corresponds to the largest amplitude motion. Arrows on **Supplementary Figures 19** and **20** show these vectors for the first principal component, if a vector (i.e. displacement) is larger than 3 Å. It can be seen that mostly the outer shell of the complexes produce large displacements and the core regions do not participate in high amplitude, correlative motions. In four out of seven complexes, at least one of the short α-helices of the monomeric units (T^75^-G^83^) is largely involved in the first component. Exceptions to this phenomenon are the (4, 3x), (4, 2x, βC) and (4, 2x, lL) complexes, where the helices (and almost all other parts of the protein) seem motionless along the first eigenmovement. In the other cases, the helices move towards or away from the axis of the lL interface. Other, highly mobile parts of the complexes are the β-turn connectors.

As the first PCA component motions seem to concentrate mostly to the outer region of the complex, a logical conclusion would be that the mutations could not alter the seemingly non-existent dynamics of the inner parts. However, we also looked at the motion patterns using RBS, which, instead of coordinate-correlations, considers atom-atom distance variations. The result of an RBS analysis is a segmentation of the protein, where each segment (atom group) can be considered relatively rigid (with respect to a threshold displacement), while larger motions are still present between these segments. **Supplementary Table 4** contains the RBS results expressed in the number of resulting segments and number of unclustered residues, while residue colorings in **Supplementary Figures 19** and **20** show how these segments are distributed in the complex. From both the numeric data, as well as from the visualizations, it is apparent that the incorporation of H88R mutations shrinks the average segment size. The WT tetramer has 4 segments with a relatively large average segment size, while the tetramer carrying the H88R mutation in all of its monomers (4, 4x) has 12 segments with a significantly smaller average segment size. Incorporation of one, two or three mutant chains seem to produce intermediate values both in the number of segments and in their sizes, outlining a strong connection between the dynamics of the complexes and the number of mutant chains inside them.

From a structural point of view, RBS segments are generally merged along the βE interface and are always divided along the lL and βC interfaces. This implies that the βE contact surface, creates a tight and rigid connection between the free β-edges, while in case of the weaker βC and lL interfaces segment-segment separation persists. We collected amino acid groupings that always appear in the same segment; this consensus segmentation can be seen in **Figure 6**, while the consensus segments (CSs) are enumerated in **Supplementary Table 5**. It is apparent that these common building blocks consist of 1, 2, 3 or 4 β-strands glued together. These can be re-merged (with the addition of non-CS residues) into the segments appearing in any of the seven RBS results. How this rebuilding happens, depends on the specific complex (**Supplementary Tables 6** and **7**). For example, CSs 3 and 10 get merged into the same segment in 5 out of 7 cases. This is not surprising, since these two CSs constitute the continuous β-sheet (containing the βE interface) between chains A and B. CSs 7 and 9 are merged in 6 out of 7 cases (they are only separated in complex (4, 3x)), while the 7, 9, 3, 10 CS composition appears in (4, 1x) and (4, 2x, βE). Similarly, for chains C and D CSs 1 and 5 are merged in all complexes, except for the complex (4, 4x), while 0, 1, 4, 5 and 6 appear together in 4 out of 7 complexes. This latter composition belongs to the continuous β-sheet between chains C and D. In complexes (4, 0x) and (4, 2x, βE), this sheet grows even further to a larger rigid segment by engulfing more CSs; 0, 1, 4, 5, 6, 8 and 11 form a single rigid body that encompasses most of (in complex (4, 2x, βE)) or all of (in complex (4, 0x), with the addition of CS 2) the β-strands in both chains.

**Figure 6.**
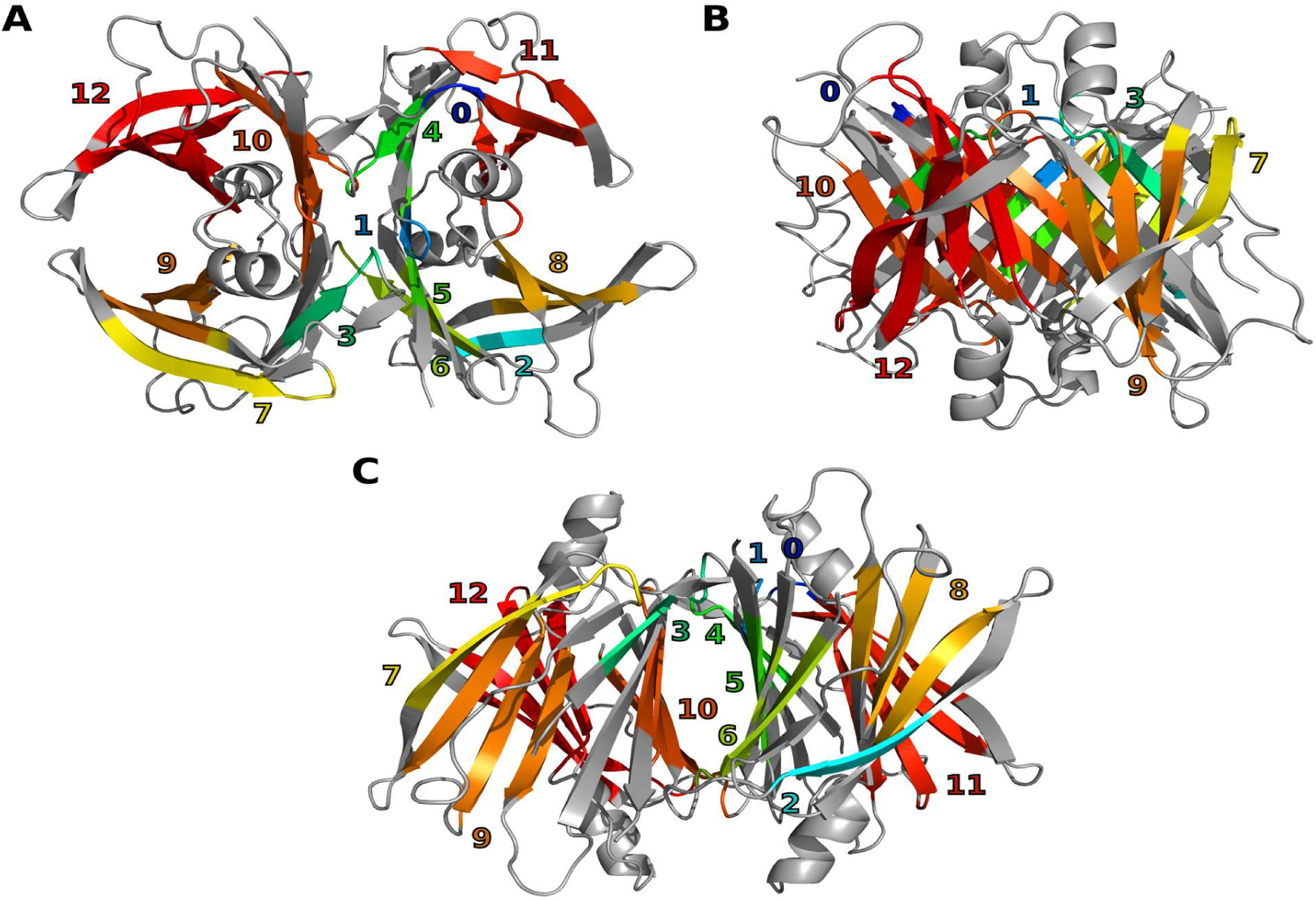
Consensus segmentation of all seven tetramers, viewed from three different angles. Residues that belong to the same segment are colored with the same color. The numbers indicate the indices of the segments. Residues that are unsegmented in any of the seven structures are colored in grey. These common segments (along with additional residues) can be considered as the basic building blocks for the actual segmentation results i.e., they can be merged and extended to give the calculated rigid body partitions for every tetramer.

## Summary

Transthyretin (TTR) amyloidosis, caused by the accumulation of transthyretin-derived amyloid depositions, is a severe, life-threatening condition, for which possible treatment options are still underexplored. The H88R variant of this protein predisposes carriers to variant transthyretin amyloidosis, necessitating a better understanding of the structure and behavior of this mutant. Here, we studied the relative stability of hybrid tetramers, which consist of both WT and H88R TTR chains in different ratios and configurations, using *in silico* molecular modeling techniques.

Using bacterial expression, WT and H88R mutant TTR proteins were successfully obtained and were mixed for a subunit exchange study. Both the size exclusion chromatography and mass spectrometry results point to the existence of hybrid tetramers. These findings have significant clinical importance to patients carrying this mutation, since the monomeric H88R chains could escape cellular housekeeping mechanisms by hijacking tetramers and can possibly get secreted to the bloodstream. Subsequent dissociation of these hybrid tetramers could then foster amyloid deposition and disease progression. In our previous study we found H88R carrier patients’ serum transthyretin levels were well below the normal range^35^ (91.3 mg/L, n=3). We argue that, in spite of hybridization, ERAD still affects H88R TTR during the asymptomatic and early stages of the disease, but it might be an important mechanism, through which, in the later stages, H88R might “leak” from hepatic cells. In spite of this, hybridization can be just as important from the therapeutic point-of-view: hybrid tetramers may be targeted by existing tetramer stabilizers. These two aspects of hybrid tetramer formation highlight a delicate balance: while TTR stabilizers can target hybrid tetramers formed by otherwise monomeric TTR variants, this stabilizing effect may also lead to increased secretion of unstable variants, as observed for A25T and D18G, where T4 binding promotes ERAD evasion and enhances secretion. Only thorough investigation of the effects of tetramer stabilizers on these species can determine the ultimate impact of this balance.

TTR variant determination of H88R mutation carrier patients supported our previous claim of decreased H88R TTR secretion. The observed low WT/H88R variant ratio, together with our computational result indicating thermodynamically unhindered hybrid tetramer formation, indicates that for probabilistic reasons H88R TTR molecules are most likely present in a 3:1 WT:H88R hybrid tetramer form after secretion. Thus, this 1x mutant hybrid tetramer may be the real target of tetramer stabilizing drugs in patients with the H88R genotype.

*In silico* studies were also performed to probe the structure, dynamics and energetics of several different hybrid and non-hybrid TTR species. In particular, 17 different 1 μs long MD simulations and 39 MM/GBSA Δ*G* calculations were performed and processed using 3 different linear mathematical models to clarify the details of complex formation energetics. The results highlight tendencies at multiple accuracy levels; the simplest model was able to order the energetic contribution of the three different interfaces, namely the βE (strongest contributor), lL and βC (weakest contributor) interfaces. The second model also considered mutation states at the interfaces, pointing out that the presence of a double H88R mutation at the βC interface is highly unfavorable, while all other single- and double H88R mutant interfaces are not significantly different from each other in formation free energies. The last model shifted energetic contributions from a relative to an absolute scale using complex-level energetic elements. This shed light to the positive Δ*G* contribution of the βC interface, a sound result considering the water-filled hydrophobic cavity at this contact surface. It also showed that the lL interface has generally no stabilizing, nor destabilizing effects to the tetramer, and the main surface holding the tetramer together is situated at the βE interface.

Local and global structural, as well as dynamical effects of the H88R mutations were also investigated in the tetramer simulations. Residue F^87^ appears as an important stabilizing hub of a hydrophobic core at the βE interface. We proved that R^88^ can occupy a wide variety of conformational niches, as shown by its χ_1_ dihedral angle distribution, solvent exposition states and contacts with the other chains. The βC and lL interfaces are also affected by the introduction of H88R mutations, although by a lesser extent, which manifests as a change in the average number of secondary chemical interactions at the contact surfaces. Dynamical effects were investigated using principal component analysis (PCA) and rigid body segmentation (RBS) analysis. PCA showed that the larges amplitude, concentrated motion modes take place on the loops connecting the β-strands of the monomers. These modes sometimes also extend towards the short α-helical part of the protein. In contrast to PCA, RBS analysis highlighted significant differences in the core regions of the different TTR hybrid tetramers. The segmentation profile indicates increased core mobility in hybrids containing more H88R chains and also smaller rigid segment sizes. Additionally, we have identified consensus segments that always appear intact in the tetrameric complexes and contain dynamically inseparable amino acids.

## Acknowledgments

Supported by the Ministry for Innovation and Technology from the Hungarian NRDI Fund (2020-1.1.6-JÖVŐ-2021-00010). Project no. 2018-1.2.1-NKP-2018-00005 has been implemented with the support provided from the National Research, Development and Innovation Fund of Hungary, financed under the 2018-1.2.1-NKP funding scheme. Project no. RRF-2.3.1-21-2022-00015 has been implemented with the support provided by the European Union. ADVANCED_25 grant (number 153035) of the National Research, Development and Innovation Fund of Hungary is also gratefully acknowledged. Project no. 2025-3.1.1-ED-2025-00013 has been implemented with the support provided from the National Research, Development and Innovation Fund of Hungary, financed under the 2025-3.1.1-ED-NKP funding scheme.

## Data Availability Statement

Code and data used for the reproduction of MM/GBSA data processing can be found at the following GitHub repository: https://github.com/fazekaszs/ttr_interface_dg.

## Author Contributions

All authors have given approval to the final version of the manuscript. Zs. F., D. K.-M. and I. B. wrote the main article text. A. P. helped with proofreading and supervised the project. I. B. and H. T.-P. performed the protein expression, purification and preparation, while D. P., D. V. and L. T. performed the mass spectrometry studies. Z. P. provided patient sera and clinical data. D. K.-M. carried out modeling of the BiP-TTR complexes. Zs. F., I. B. and H. T.-P. prepared the *in silico* molecular systems and started the molecular dynamics runs. Zs. F. and H. T.-P. performed downstream *in silico* studies. A. P. served funding for the project.

